# Oncogenic Role of Aberrant EZH2 in Hepatoblastoma

**DOI:** 10.1101/2025.07.30.667506

**Authors:** Kathryn Glaser, Lara Berklite, Brian T. Hickner, Priyanka Rao, Roma H. Patel, Andrew A. Badachhape, Somak Roy, Emily Schepers, Nikolai A. Timchenko, James I. Geller, Sarangarajan Ranganathan, Gregory M. Tiao, Bruce Aronow, Takanori Takebe, Sarah E. Woodfield, Sanjeev A. Vasudevan, Erica A.K. DePasquale, Alexander J. Bondoc

## Abstract

Hepatoblastoma (HB) is the most common pediatric liver malignancy. However, its cellular origin and molecular drivers remain poorly defined. Using single-nuclear RNA sequencing (snRNA-seq), we identified a proliferative, hepatocyte-derived tumor cell population (cycling Hep^T^) enriched for Enhancer of Zeste Homolog 2 (*EZH2)* expression, particularly in the aggressive embryonal subtype. Integrative genomic and transcriptomic profiling confirmed *EZH2* overexpression. Disruption of the PRC2 complex was evident through mislocalization and reduced expression of SUZ12, a core component. *EZH2* overexpression correlated with upregulation of mitotic regulators such as *AURKB* and *Ki67* in human HB gene expression analysis as compared to background liver. Targeted sequencing identified variants of uncertain significance in *EZH2* and *SUZ12* in 11 of 11 patient tumors. Pharmacologic inhibition of EZH2 with EPZ-6438 reduced proliferation and sensitized HB cells to cisplatin through gene regulation, potentially modulating platinum accumulation both *in vitro* and *in vivo*. In summary, EZH2 promotes HB progression through epigenetic silencing and noncanonical signaling pathways. These findings support EZH2’s contribution to HB pathogenesis, therefore identifying it as a novel therapeutic target.

**STATEMENT OF SIGNIFICANCE:** EZH2 is enriched in hepatoblastoma, suggesting its role in early tumorigenesis. Its inhibition, combined with cisplatin, produces synergistic anti-tumor effects, supporting EZH2 as both an oncogenic and therapeutic target.

## INTRODUCTION

Hepatoblastoma (HB), the most common primary hepatic malignancy in children, is a rare and aggressive form of liver cancer. Therapeutic options for advanced disease combine neoadjuvant chemotherapy and surgical resection, sometimes including liver transplantation, which together have potential long-term complications such as the risk of rejection, relapse, and ototoxicity (1-3). The clinical histopathology of HB reveals a heterogeneous tumor composed of various cell types, including epithelial (fetal, embryonal, and mixed histology) and mesenchymal elements. Histopathological examination is crucial for diagnosis, therapeutic stratification, and prognosis (4). The presence of small, round blue cells, fetal hepatocytes, and embryonal components are characteristic features of HB (4). Overexpression of the epigenetic regulator *enhancer of zeste homolog 2* (*EZH2*) contributes to HB tumor growth and metastasis (5,6). This study explored the mechanism by which EZH2 overexpression drives the proliferation of a novel mitotic cell population critical to HB pathogenesis and explored the therapeutic potential of EZH2 inhibition.

EZH2 is a histone methyltransferase that catalyzes the trimethylation of histone H3 at lysine 27 (H3K27me^3^), leading to epigenetic silencing of target genes. This epigenetic modification has been linked to the regulation of various cellular processes, including cell proliferation, differentiation, and migration (7). EZH2 overexpression in HB is associated with increased tumor cell proliferation, invasion, and metastasis. It is unclear whether *EZH2* acts through epigenetic regulation, noncanonical mechanisms, or possibly through a dysregulated interplay between the two (8-10).

Oncologically, canonical *EZH2* overexpression upregulates cell-cycle progression genes including cyclins and cyclin-dependent kinases leading to increased rates of cell division and higher mitotic indices within tumor cell populations (11). Furthermore, EZH2 modulates the activity of other key cell cycle regulators, such as Rb protein and E2F transcription factors thereby disrupting cell cycle checkpoints (12,13). Additionally, EZH2 maintains a stem-like cell population within the tumor, which is thought to drive tumor progression and metastasis (14,15). EZH2 is upregulated in a subset of HB cells that exhibit increased metastatic potential (16).

Alternatively, noncanonical EZH2 signaling pathways have also been implicated in the regulation of mitotic processes in HB cells. For example, EZH2 interacts with the Aurora kinase family, which are critical regulators of mitotic spindle formation and chromosome segregation (13). This interaction can lead to the dysregulation of these mitotic processes, contributing to chromosome instability and the accumulation of genetic aberrations within tumor cells (17,18). The heterogeneous cellular nature of HB with subpopulations demonstrating varying degrees of EZH2 overexpression, may contribute to the development of treatment resistance and disease recurrence (19). Given EZH2’s possible role promoting a more aggressive HB phenotype, inhibition of this pathway may yield novel therapeutic strategies.

In this study, we explore the molecular landscape of HB by analyzing distinct tumor cell clusters with a particular focus on one characterized by *EZH2* overexpression termed “cycling Hep_T_.” We also evaluated HB response to EZH2 inhibition in both *in vitro* and *in vivo* models. For *in vitro* studies, HB cell lines and patient tumor-derived cells were compared to other liver cancer controls including hepatocellular carcinoma (HCC) and hepatocellular neoplasm not otherwise specified (HCN-NOS). EZH2 is markedly overexpressed in HB and drives proliferative signaling, genomic instability, and maintenance of metastatic subpopulations. Its dual function in PRC2-mediated epigenetic repression and noncanonical oncogenic pathways underscores its role as a driver of tumor aggressiveness.

## MATERIALS AND METHODS

### Human tissue specimens

Human background liver and tumor samples were collected from patients with HB and HCC under conditions appropriate for characterization and generation of cell models with institutional review board approval (IRB # 2016-9497, 2023-0451, PI-Bondoc at Cincinnati Children’s Hospital Medical Center) and informed patient consent. Data from 56 patient tumors (**Supplemental Table S1**) and cell models from three HB patient samples are reported across all methodologies.

### Immunohistochemistry and pathology review

Formalin-fixed, paraffin-embedded tumor samples were used for immunohistochemical analysis of EZH2 expression. Immunohistochemistry (IHC) staining was performed on paraffin sections for hematoxylin and eosin (H&E) and EZH2 (3147, 1:200; Cell Signaling Technologies). Tissue sections were deparaffinized and rehydrated, and the antibody was diluted in phosphate-buffered saline (PBS)-containing 2% BSA and 0.05% Tween 20. Tissue sections were incubated with antibody overnight at 4°C, washed with PBS + 0.05% Tween 20 and then visualized with Vectastain elite ABC HRP (Vector Labs) per protocol instructions. Serial sections were imaged. Staining intensity and proportion of positive tumor cells were assessed by a pathologist who was blinded to the clinical outcome. Intensity was graded as 1+/2+/3+ and percentage of tumor cells showing positive staining at each intensity level was recorded for each tumor. Intensity of staining in each morphologic pattern (i.e. fetal, embryonal, etc), if more than one was present in the tumor, was also graded using the same scale. H&E sections for each tumor were reviewed to identify the morphologic patterns present in each case.

### Immunofluorescence

The cells were grown on chamber slides and fixed with ice-cold methanol. Cells were washed with 1x PBS, blocked with 2% BSA and 0.05% Tween 20, and incubated with primary antibody overnight at 4°C followed by incubation with secondary antibody for 1 hour at room temperature (Cell Signaling antibodies were used: EZH2 3147, 1:200, H3K27me3 9733, 1:1,000, EED 51673, 1:200, SUZ12 3737, 1:200).

### CinCseq mutation Panel

To assess the mutational landscape of HB samples, we performed targeted next-generation sequencing using a custom panel of cancer-related genes. For formalin-fixed paraffin-embedded (FFPE) specimens, Hematoxylin & Eosin (H&E) stained slides were reviewed by a pathologist and the target area(s) for sequencing was marked. Manual macro- or microdissection of the targeted tumor region was performed using unstained tissue sections under H&E guidance. Total nucleic acid (TNA) from whole blood and bone marrow samples and DNA and RNA from FFPE specimens were isolated using standard laboratory procedures. After fragmentation and library preparation using xGen™ cfDNA & FFPE DNA Library Preparation Kit (IDT, Coralville, Iowa), short DNA/cDNA fragments were enriched using custom hybrid capture probes (IDT, Coralville, Iowa/GOAL consortium) and sequenced using a 2 × 150bp sequencing chemistry on an Illumina 2000. Clinically validated CinCSeq NGS analysis evaluates single nucleotide variants (SNV), multi-nucleotide variants (MNV), and insertions and deletions (indels) in approximately 95% of the DNA coding exon sequences of 383 cancer-related genes, copy number variants (CNV) in 531 cancer-related genes, and gene fusions/structural variants in 183 cancer-related RNA genes.

A custom bioinformatic analysis pipeline for DNA and RNA sequencing was performed using the GRCh38/hg38 human reference genome (GCA_000001405.29). Briefly, bcl convert (Illumina) was used to demultiplex and generate FASTQ files. After pre-alignment processing using fastp (20), the sequencing reads in the FASTQ files were aligned against the human reference genome using bwa-mem2(21) and STAR aligner (22) for DNAseq and RNAseq data, respectively. For DNAseq, after sorting and indexing using samtools (23), deduplication using sambamba (24), and indel realignment using Abra2 (25), variants were identified using VarDict (26) and FLT3-ITD-ext (27). Called variants were normalized using bcftools (23) and annotated using hgvs python package (28) and echtvar (29).

The analytic sensitivity is 3-5% variant allele fraction (VAF) for the detection of SNVs, MNVs, and small Indels (<10bp) and 5-10% VAF for larger indels (>10bp) and structural variants, including internal tandem duplication (ITD) variants. The minimum required sequencing depth of coverage is 300x. Orthogonal confirmation for one or more genomic alterations may be performed if appropriate. Interpretation of genomic alterations was performed following the AMP/ASCO/CAP guidelines (30) for the interpretation of somatic variants. The HUGO gene nomenclature and HGVS variant nomenclature systems were used for reporting sequence variant information. Tumor mutational burden (TMB) was determined by measuring the number of non-synonymous single nucleotide variants in the coding region of the sequenced genes on the CinCSeq test that include AMP/ASCO/CAP tier 1/2/3 variants which are at 3% VAF or higher after filtering. TMB was reported as the number of mutations per megabase (Muts/Mb).

### Single Nuclear RNA Sequencing

Nuclei from human tumor and liver were extracted, libraries were constructed, and sequenced by 3V3 10X Genomics, as described previously (16). Cell Ranger software (version 6.0.0) was used to generate count matrices for each sample following standard practices(31). SoupX was then used to correct for ambient RNAs in each sample using auto-estimation of the contamination fraction (32). Quality control was performed on each sample by retaining only those cells with 1000 or more UMIs, 500 or more features, and less than 20% mitochondrial fraction. This resulted in 110,257 cells across the 11 HB tumors, six background liver, and one non-malignant control samples. Seurat version 5.1.0 was used to log-transform and scale data following developer recommendations (33,34). All samples were integrated with Seurat and the integrated dataset was used for principal component analysis and cluster identification (35).

### Cell Classification

After initial clustering at a resolution of 1, which resulted in 24 clusters, sub-clustering was performed to define biologically relevant cell types. The expression of known marker genes and gene sets, pathway analysis, and differential expression analyses were used to aid in identification, which resulted in 11 distinct broad cell types. Proportional tables and bar plots were generated to examine the compositional differences between background and tumor cell populations, which were statistically validated using a Wilcoxon test. Dot plots were generated using Seurat scaling to visualize the key marker genes used in cluster identification. Cells were scored for cell cycle gene expression using the S and G2/M phase gene sets loaded with Seurat, and scores were overlaid onto subcluster UMAPs within Seurat.

### Gene Set Enrichment Analysis (GSEA)

Pre-ranked GSEA was used to identify regulatory pathways enriched in the differential expression comparison between tumor and background per cell type (36,37). The Human Molecular Signatures Database (MSigDB) Hallmark gene sets and Transcription Factor Targets regulatory gene sets were used for scoring (38,39). Gene sets were considered enriched for this analysis if their false discovery rate was less than 25% and if their nominal p-value was less than 1%, in line with GSEA’s standard reporting.

### Cell Culture

We used established HB cell lines and patient-derived HB cell models to assess the functional consequences of EZH2 inhibition and subsequent chemotherapy treatment. The cells were grown in DMEM supplemented with 10% FBS, 1% NEAA, and 1% penicillin/streptomycin. *In vitro* cells were treated with EZH2 inhibitors (tazemetostat (EPZ-6438) 10μM-Selleck, GSK126 5μM, Tocris) for 72 hours. Cisplatin (0-400μM) was then administered for 24 hours. HepG2 cells (passage 5-25) were purchased from ATCC and not authenticated. Additional commercially available cell lines, HUH6 and HUH7 (passage 5-25), were kindly shared by the Timchenko laboratory. HBc108, 129, and 130 (passage 4-10) were derived from HB patient tumor by explant culture in the media listed above plus 20 μM Rho kinase inhibitor, Y27632. All cell lines were routinely monitored for mycoplasma and tested negative using a Mycoalert detection kit (Lonza).

### qRT-PCR

Total RNA was extracted using the RNeasy Plus Mini Kit (Qiagen), and cDNA was synthesized using SuperScript VILO (Invitrogen). Quantitative PCR (qPCR) was performed using SYBR Green Master Mix (Qiagen) on a CFX96 Real-Time PCR System. RT^2^ qPCR primers were used (Qiagen). The relative expression was calculated using the ΔΔCt method, normalized to GAPDH (**Supplemental Table S2**).

### Patient Derived Xenograft (PDX) Model and Treatment Protocol

Murine studies were conducted following ethical and compliance standards and with Institutional Animal Care and Use Committee (IACUC) approval (protocol AN-6191, PI-Vasudevan at Baylor College of Medicine). Tissues for PDX models were obtained from patients consented to IRB protocol H-38834 (PI-Vasudevan at Baylor College of Medicine). Tumor tissue fragments (3-5 mm^3^) were implanted orthotopically into the livers of 6-12-week-old NOD scid gamma (NSG) immunocompromised mice, as previously described (40). Two weeks post-implantation, tumor growth was monitored using magnetic resonance imaging (MRI). To better model the tumor burden observed in patients with advanced HB, treatment was initiated once tumor volumes reached 0.1 cm^3^ or greater, as confirmed by MRI.

Mice were treated with cisplatin at a dose of 5 mg/kg administered once weekly via intraperitoneal injection and/or EPZ-6438 (Selleck) at a dose of 150 mg/kg administered five times per week via oral gavage. The control mice received no treatment. Treatments were continued for approximately 4 weeks or until mice reached a humane endpoint, defined as a tumor diameter of 1.5 cm. Of note, two mice treated with cisplatin alone were euthanized early, after one week of treatment, as part of another study protocol conducted simultaneously. EPZ-6438 was prepared according to the manufacturer’s guidelines, suspended at a concentration of 15mg/ml in 0.5% CMC-Na and 0.1% Tween. Cisplatin was diluted in sterile saline. Tumor response was assessed through serial tumor volume calculation, which was obtained using MRI imaging. The mice appeared healthy without any weight loss throughout the treatment period.

### Statistical Analysis

Differential cell type proportion analysis was performed using the two-tailed Wilcoxon test. Differential expression analysis was performed using FindAllMarkers and FindMarkers within Seurat version 5.1.0 using the default parameters for the Wilcoxon test. Additionally, gene expression, mitotic cell number, and cell viability data were analyzed using GraphPad Prism software. Statistical significance was determined using paired Student’s t-test (Tumor/Liver gene expression) or ANOVA with multiple comparisons (subtype comparisons, drug treatment, and mitotic cells) as appropriate. Statistical significance was set at P < 0.05.

## Data Availability

The data generated in this study are publicly available in the Gene Expression Omnibus (GEO) repository under accession number GEOXXXXXXX. The code used to analyze the data and generate the figures used in this study can be found at the project GitHub: https://github.com/EDePasquale/Bondoc-HB.

## RESULTS

### Single-Nuclear RNA Sequencing Defines HB Tumor Cell Subtypes

Single nuclear RNA sequencing (snRNA-seq) was used to profile HB tumor transcriptomes, identify key cell types, marker genes, and dysregulated pathways for further analysis. The integrated single nuclear RNA sequencing (snRNA-seq) dataset was first examined to ensure there were no major sample-specific clusters or other batch effects that could skew downstream analyses. Projecting sample identities onto the unified UMAP plot (**Figure 1A**) reveals no clusters dominated by individual samples or a limited subset of samples, indicating successful integration. A combination of known marker genes for liver cell types and pathway analysis was used to define 11 distinct cell type populations (**Figure 1B**), which were prevalent in both background and tumor samples (**Figure 1C**). The broad hepatocyte cluster showed the most spatial discrimination between samples within the UMAP, which is likely due to high diversity between hepatoblast tumors in this cell type and not technical artifacts of integration.

**Figure 1.**
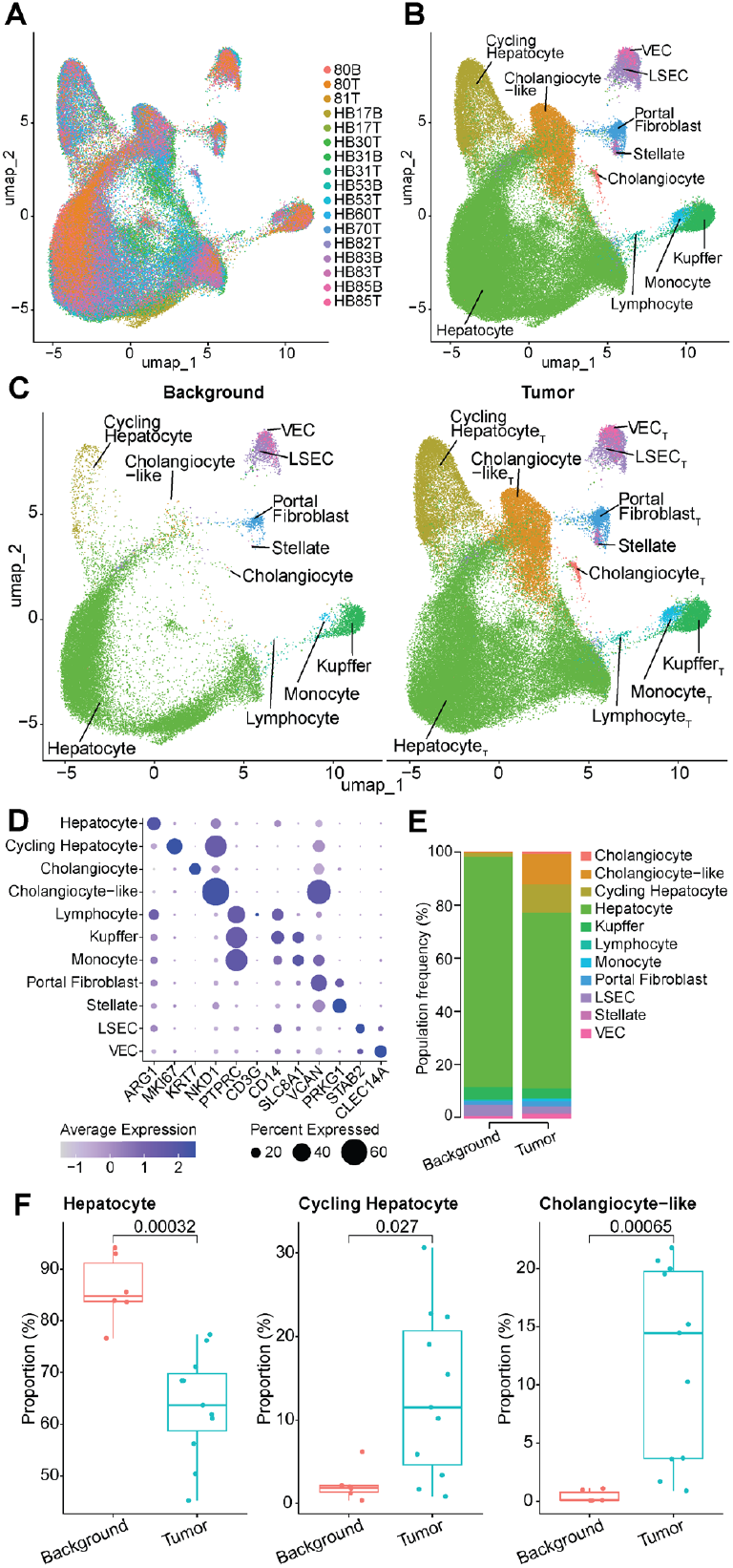
Integration and annotation of cellular subpopulations in tumor and background HB samples. A. UMAP plot showing successful integration of 17 background and tumor samples, colored by sample ID. B. UMAP colored by 11 broad cell types. C. Same UMAP as panel B but split to show only background (left) and tumor (right) samples. D. Dot plot showing average normalized expression (color) and the percent of cells expressing the gene (dot size) for established lineage and cluster markers. E. Stacked proportional bar plot showing the prevalence of each cluster within the background (left) and tumor (right) samples. All clusters add up to 100% for each condition. F. Box plots for the Hepatocyte (left), Cycling Hepatocyte (middle), and Cholangiocyte-like (right) clusters showing decreased prevalence in tumor for Hepatocyte (p = 0.00032), and increased prevalence in tumor for Cycling Hepatocyte (p = 0.027) and Cholangiocyte-like (p = 0.00065) clusters. A Wilcoxon test was used for statistical analyses.

For each of these broad cell types, we defined key markers for identification, which we visualized here as a dot plot (**Figure 1D**). Epithelial populations were defined by *ARG1* for hepatocytes and *KRT7* for cholangiocytes. The cycling Hep_T_ population was evident from *MKI67* expression. *NKD1* was strongly expressed in the cholangiocyte-like population or hybrid tumor cell with biliary and hepatocyte marker genes as well as the cycling hepatocytes. The three immune populations were all defined by expression of *PTPRC*, with *CD3G* expression identifying lymphocytes, *CD14* and *SLC8A1* expression identifying Kupffer cells, and *SLC8A*1, and *VCAN* identifying monocytes. Mesenchymal populations were defined by higher levels of *VCAN* expression in portal fibroblasts and *PRKG1* expression in stellate cells. Finally, liver sinusoidal endothelial cells (LSECs) were identified by high *STAB2*, and vascular endothelial cells (VECs) were defined by *CLEC14A* expression.

Differences in cell type proportions observed in the UMAP were further visualized using a proportional stacked bar plot (**Figure 1E**). We found that the background population was dominated by hepatocytes, while the tumor population had a relatively higher proportion of other cell types. Proportional differences for the three larger populations were statistically significantly different via Wilcoxon test (p < 0.05) between background and tumor conditions (**Figure 1F**), with background samples having proportionally more hepatocytes (p = 0.00032) and tumor cells having more cycling hepatocytes (p = 0.027) and cholangiocyte-like cells (p = 0.00065).

### EZH2 Dysregulation in Tumor-Specific Hepatocytes

We next focused on the main tumor “hepatocyte” populations including the hepatocyte, cycling Hep_T_, and cholangiocyte-like populations as compared to those in background liver samples, which were subset from the larger UMAP (**Figure 2A**). As seen in the previous proportional bar plot, the background samples were primarily comprised of traditional hepatocytes, whereas the tumor samples contain relatively more cycling hepatocytes and cholangiocyte-like hepatocytes (**Figure 2B**).

**Figure 2.**
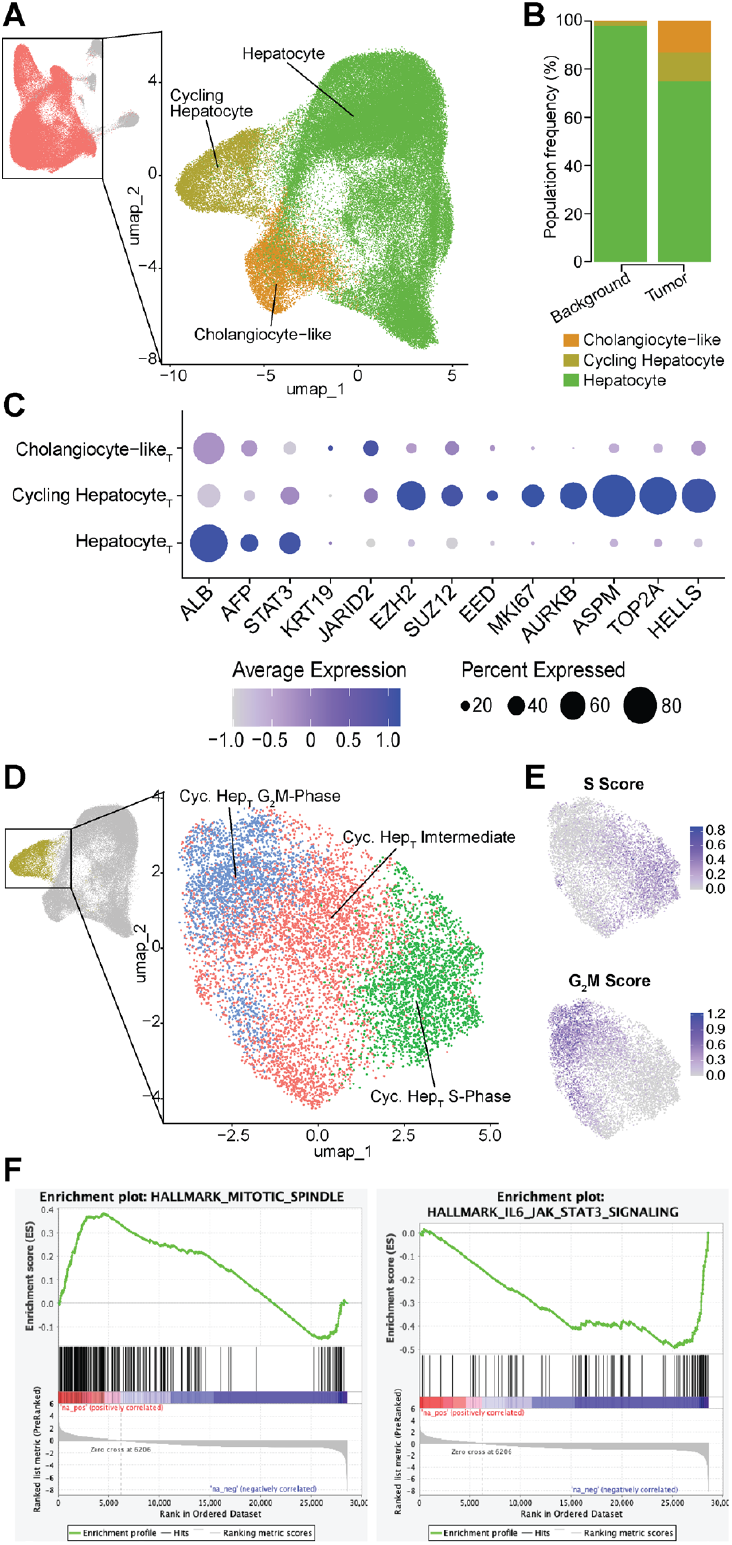
Dissecting HB tumor heterogeneity through subclustering and functional pathway analysis. A. UMAP showing the Hepatocyte sub-clustering, with the upper-left UMAP colored in salmon to show which cells were selected for sub-clustering, and the larger right-hand UMAP showing the three sub-clusters that were derived using unsupervised clustering. B. Stacked proportional bar plot showing the prevalence of each Hepatocyte sub-cluster in background (left) and tumor (right) samples. All clusters add up to 100% for each condition. C. Dot plot showing average normalized expression (color) and the percentage of cells expressing the gene (dot size) for marker genes used to define the Hepatocyte sub-clusters. D. UMAP showing Cycling Hepatocyte sub-clustering, with the upper-left UMAP colored in gold to show which Hepatocyte sub-clusters were selected for further sub-clustering, and the larger right-hand UMAP showing the three resultant Cycling Hepatocyte sub-clusters. E. The same Cycling Hepatocyte sub-cluster UMAP plot colored by Seurat-derived S Score (top) and G_2_M Score (bottom), with darker blue indicating higher expression. F. Gene Set Enrichment Analysis (GSEA) plots showing a positive enrichment score in tumor cells compared to background for the ‘HALLMARK_MITOTIC_SPINDLE’ gene set and a negative enrichment score for the ‘HALLMARK_IL6_JAK_STAT3_SIGNALING’ gene set.

When specificially examining the tumor samples alone, we found that the cycling Hep_T_ population expressed high levels of *EZH2, SUZ12*, and *EED*, along with cell cycle regulators *MKI67, AURKB, ASPM, TOP2A*, and *HELLS* (**Figure 2C**). The tumor hepatocytes expressed high levels of *ALB, AFP*, and *STAT3*, with the later two genes known to be altered in liver cancer (4) and fibrosis (41), respectively. *STAT3* can also be repressed by *EZH2* through canonical epigenetic mechanisms (8). When compared to other published studies these integrated tumor populations are consistent with clusters defined as hepatocytic and progenitor populations as well as with greater presence in embryonal tumors (16,42,43).

The tumor cholangiocyte-like hepatocytes expressed *KRT19* at higher levels than hepatocytes, as well as *JARID2*, which has been shown to be upregulated in liver cancers (44). Further examination of the cycling Hep_T_ population in the tumor samples revealed three sub-clusters defined by cell cycle stage: Cycling Hepatocyte G_2_M-Phase, Cycling Hepatocyte S-Phase, and an intermediate population termed Cycling Hepatocyte Intermediate (**Figure 2D**). Seurat scoring for cell cycle phase informed the naming of these clusters (**Figure 2E**).

Gene set enrichment analysis (GSEA) of the cycling Hep_T_ cluster revealed significant enrichment for pathways involved in mitotic spindle assembly, while JAK-STAT signaling was notably downregulated. Elevated mitosis aligns with the enhanced replication and cell cycle signature of the cycling Hep_T_ cluster. The reduction in JAK-STAT activity may reflect epigenetic repression mechanisms, potentially mediated by *EZH2*, and aligns with our broader findings of altered *STAT3* regulation in HB. (**Figure 2F**).

### EZH2 Overexpression Correlates with Embryonal Histologic Subtype

HB and HCC histologic samples were blinded and reviewed by clinical pathologists (Berklite, Ranganathan) to define the histologic pattern as well as score the expression of EZH2 immunostaining. HCC specimens were utilized as a disease control against HB, given the differential clinical characteristics of pediatric HCC (e.g., affecting older children and more aggressive clinical behavior).

Although the EZH2 staining pattern was somewhat variable from sample to sample, EZH2 protein expression was generally highest in the embryonal regions of HB sections (**Table 1, Figure 3A**). In comparison, EZH2 staining in HCC samples was highly variable. Additional staining showed elevated expression of EZH2 across different subtypes of HB, including mixed epithelial histology, embryonal (E), and crowded fetal (CF). Elevation of SUZ12 was histologically inconsistent, whereas EED expression was observed across tumor subtypes, as well as in normal liver (**Figure 3A**). While some tumors showed coordinated upregulation of all three components, others displayed discordant expression patterns, indicating heterogeneity in PRC2 complex composition and potential functional divergence across tumor subtypes.

**Table 1.**
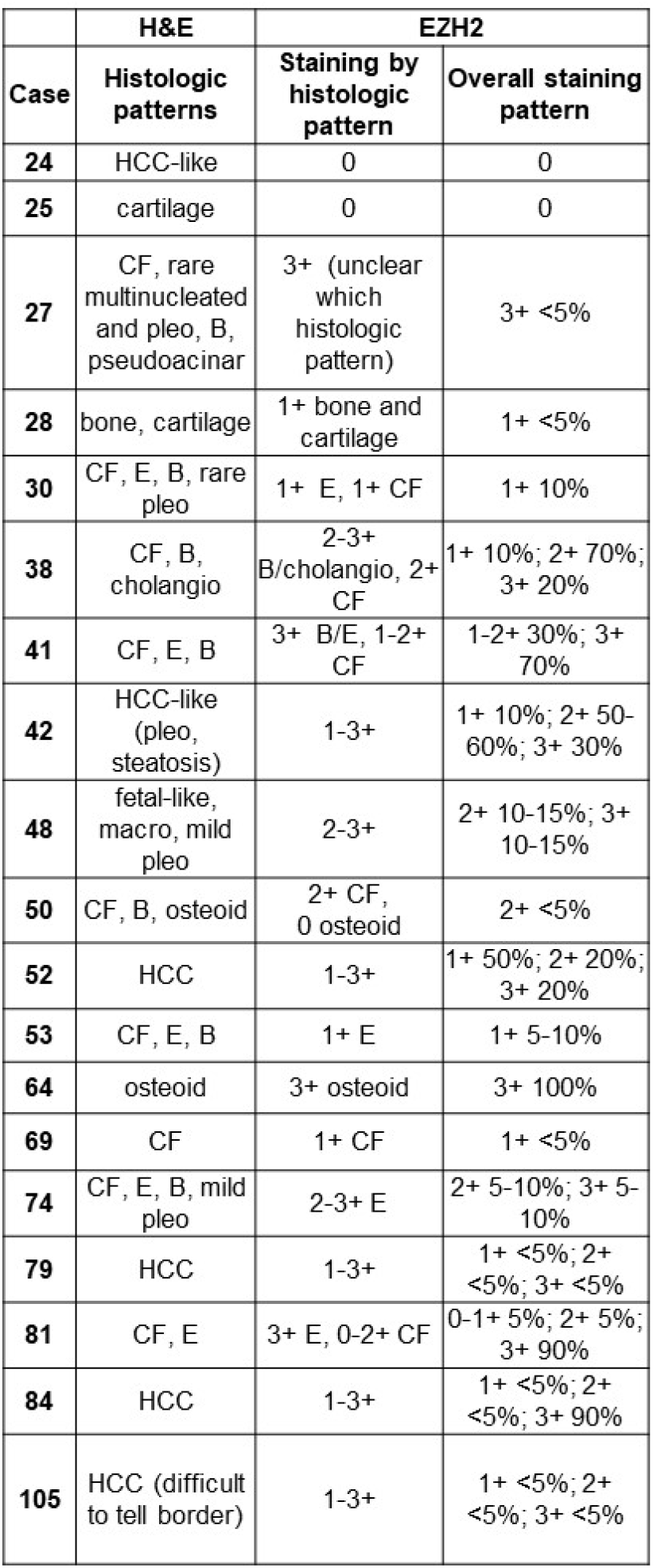
EZH2 staining pattern in HB relative to histopathologic subtype in tumor. EZH2 is overall more prevalent in embryonal histology. Extreme variability is observed in staining of fetal and mixed epithelial histology. HCC generally has less EZH2 staining. Key: CF-crowded fetal, E-embryonal, B-blastemal, macro-macrotrabecular, cholangio-cholangioblastic, pleo-pleomorphic

**Figure 3.**
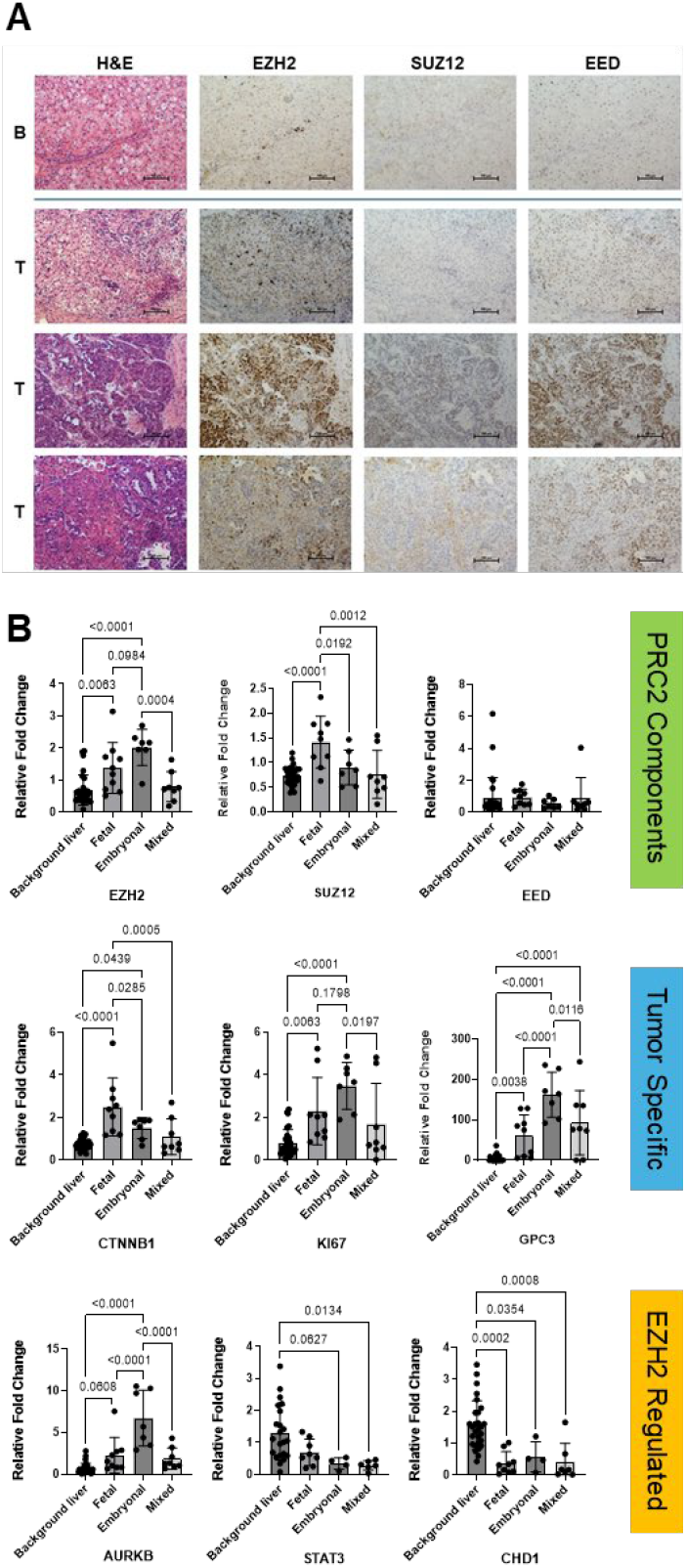
Tumors with embryonal histology have features of cycling Hep_T_ cell cluster. A. EZH2 is elevated in HB tumor though to varying degree based on histologic subtype. PRC2 components, SUZ12 and EED are often upregulated compared to normal adjacent liver, but protein expression patterns are not consistent. B. Gene expression was analyzed in tumor grouped by histologic subtype to look at PRC2 genes, tumor markers, and genes regulated by EZH2. EZH2 gene expression is highest in embryonal subtypes, as well as AURKB, Ki67, and GPC3. Fetal histology had the highest level of SUZ12 and CTNNB1 expression, where EED gene expression did not show elevation by subtype. Tumors with mixed epithelial histology had muted expression levels making interpretation difficult.

Transcriptomic profiling of 25 HB tumors segregated by clinical histopathologic subtype revealed that *EZH2* expression was elevated in HB samples with embryonal and fetal histology compared to normal liver tissue, with the highest levels observed in tumors with embryonal histology (**Figure 3B**). Surprisingly, *SUZ12* gene expression showed the highest upregulation in fetal histology with no significant difference seen in embryonal histology compared to normal liver, and *EED* gene expression was not significantly higher than normal liver regardless of subtype. This suggests that baseline expression is sufficient for canonical regulation or that other regulatory mechanisms may also be at play.

In addition to PRC2 components, we explored known HB tumor-associated genes to correlate with *EZH2* expression patterns by tumor subtype. *GPC3* and *Ki67* gene expression patterns correlate with *EZH2* expression with higher expression in embryonal tumors. *CTNNB1* (beta-catenin) showed the highest expression in predominantly fetal tumors but in embryonal tumors as well. Overall, expression in tumors with mixed histology was variable. Genes potentially regulated by *EZH2* were also explored. *AURKB*, which was previously identified in the same HB tumor cell subpopulation as *EZH2* (16), was consistently elevated along with *EZH2* across subtypes. *STAT3* expression was markedly reduced in both embryonal and mixed histology subtypes, contrasting with its typical oncogenic role. *CDH1* was also significantly downregulated in embryonal and fetal HB, suggesting the potential for promoting EMT and oncogenesis.

### *EZH2* Overexpression Does Not Consistently Correlate with other PRC2 Components

Quantitative PCR analysis revealed significant alterations in gene expression between benign (B) and tumor (T) tissues. *EZH2, AURKB, GPC3, CTNNB1*, and *TGFβ* were significantly upregulated in tumor tissues, indicating their potential roles in tumor progression (**Figure 4A**). In contrast, *MYC, STAT3, CDH1*, and *CCND1* were significantly downregulated in tumor tissues compared to normal controls, suggesting a loss of tumor-suppressive or differentiation-related functions. Although *SUZ12* and *EED* showed trends toward increased expression, these changes did not reach statistical significance (p = 0.0573 and p = 0.2123, respectively).

**Figure 4.**
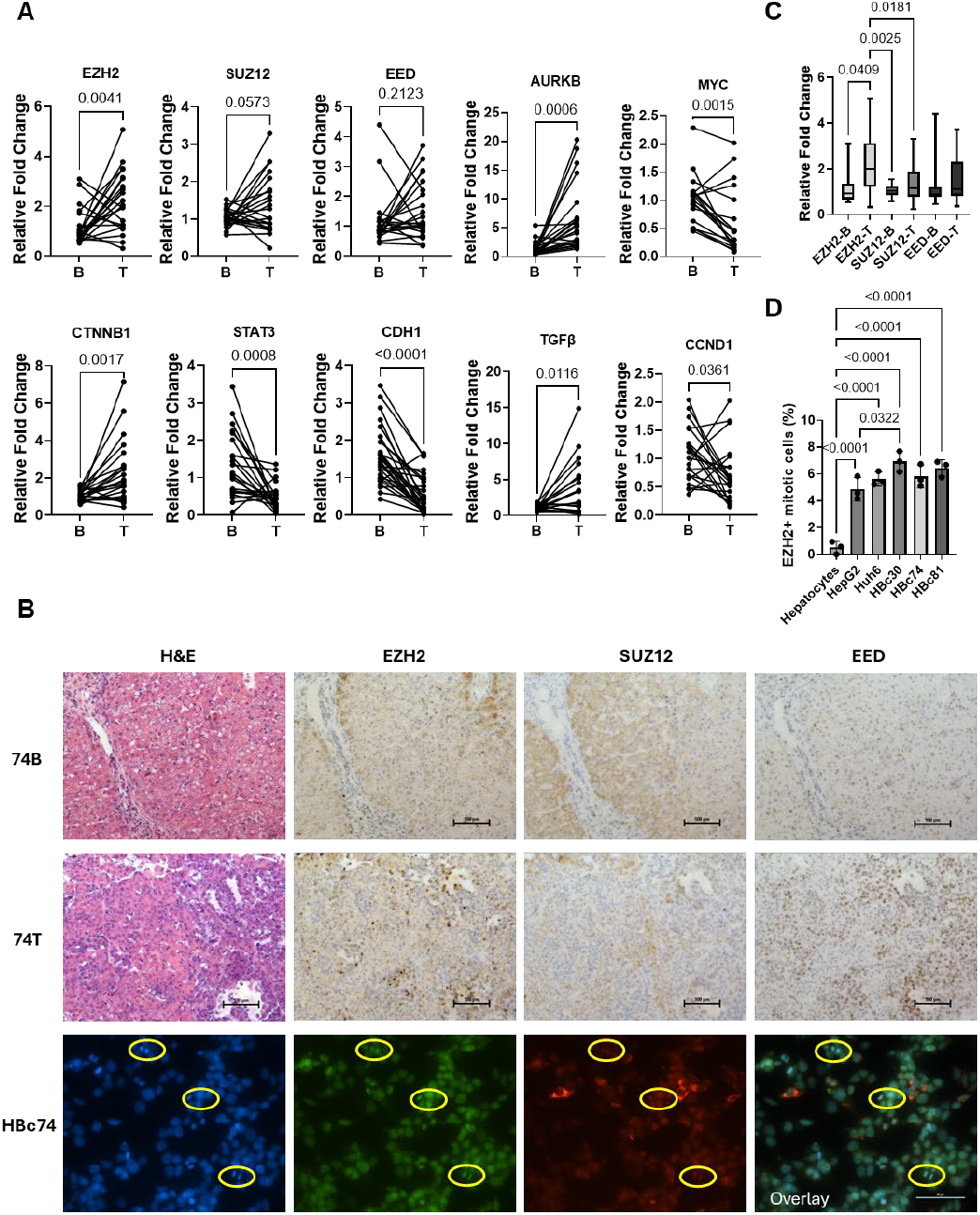
Complex regulation through canonical and noncanonical mechanisms. A. Gene expression analysis of paired liver and HB tumor samples demonstrated upregulation of EZH2, AURKB, CTNNB1, and TGFβ. EED and SUZ12 are also upregulated though not significantly. MYC, CCND1, STAT3, and CDH1 are downregulated in tumor compared to liver. B. EZH2 protein expression in tumor is not in stoichiometric relationship with other PRC2 components. EZH2 and EED are observed in the nucleus. EZH2 expression is also observed in mitotic cells from HB tumor, where SUZ12 expression is highest in the cytoplasm. C. Paired patient liver and HB tumor gene expression was compared and demonstrated significant upregulation of EZH2 in HB. Slight upregulation was also seen in PRC2 components, SUZ12 and EED, with correlation between these genes in paired samples but inconsistent upregulation. D. EZH2 positive, mitotic cells are significantly increased in HB cells compared to normal human hepatocytes. This expression pattern is also elevated in HB patient derived cells compared to HepG2 cells, though not all HB cells show a statistical significance increase compared to HepG2.

Histological and immunohistochemical analyses supported these findings. Hematoxylin and eosin (H&E) staining revealed increased cellular density in tumor samples, while immunohistochemistry confirmed elevated EZH2 protein expression in tumor tissues (74T) relative to benign tissues (74B) (**Figure 4B, C**). However, SUZ12 and EED staining did not consistently show increased expression across samples.

Fluorescence microscopy further examined the subcellular localization of PRC2 components. EZH2 and EED were predominantly localized to the nucleus in tumor cells, consistent with their known roles in chromatin modification. In contrast, SUZ12 did not co-localize with EZH2 or EED, suggesting a possible disruption of PRC2 complex formation or altered subcellular dynamics in the tumor context. This inconsistency was also observed at the gene expression level. The analysis of mitotic cell populations revealed a significant increase in the percentage of EZH2-positive mitotic cells in tumor tissues compared to benign controls (**Figure 4D**).

### EPZ-6438 and GSK-126 Enhance Cisplatin Sensitivity in Liver Cancer Cell Lines

To evaluate the effect of EZH2 inhibition on cisplatin sensitivity, six liver cancer cell lines (HBC108, HBC129, HBC130, HepG2, HUH7, and HUH6) were treated with increasing concentrations of cisplatin (0–400 µM), either alone or in combination with EPZ-6438 (10 µM) or GSK-126 (5 µM).

Cisplatin monotherapy induced a dose-dependent reduction in cell viability across all cell lines, though with reduced sensitivity in some samples (**Figure 5A**). Combination treatments with EPZ-6438 or GSK-126 significantly enhanced the cytotoxic effect of cisplatin, particularly at higher concentrations (100–400 µM). The most pronounced synergistic effects were observed in HUH6 and HBc130 cells, where combination treatments resulted in significantly lower viability compared to cisplatin alone. Statistical analysis confirmed that both inhibitors significantly sensitized cells to cisplatin, with GSK-126 showing slightly greater efficacy in some cell lines. Morphological evidence of enhanced cytotoxicity was also observed microscopically in HepG2 and HUH6 cells with the most pronounced reduction in viability observed in HUH6 combination treatment conditions further supporting the quantitative findings.

**Figure 5.**
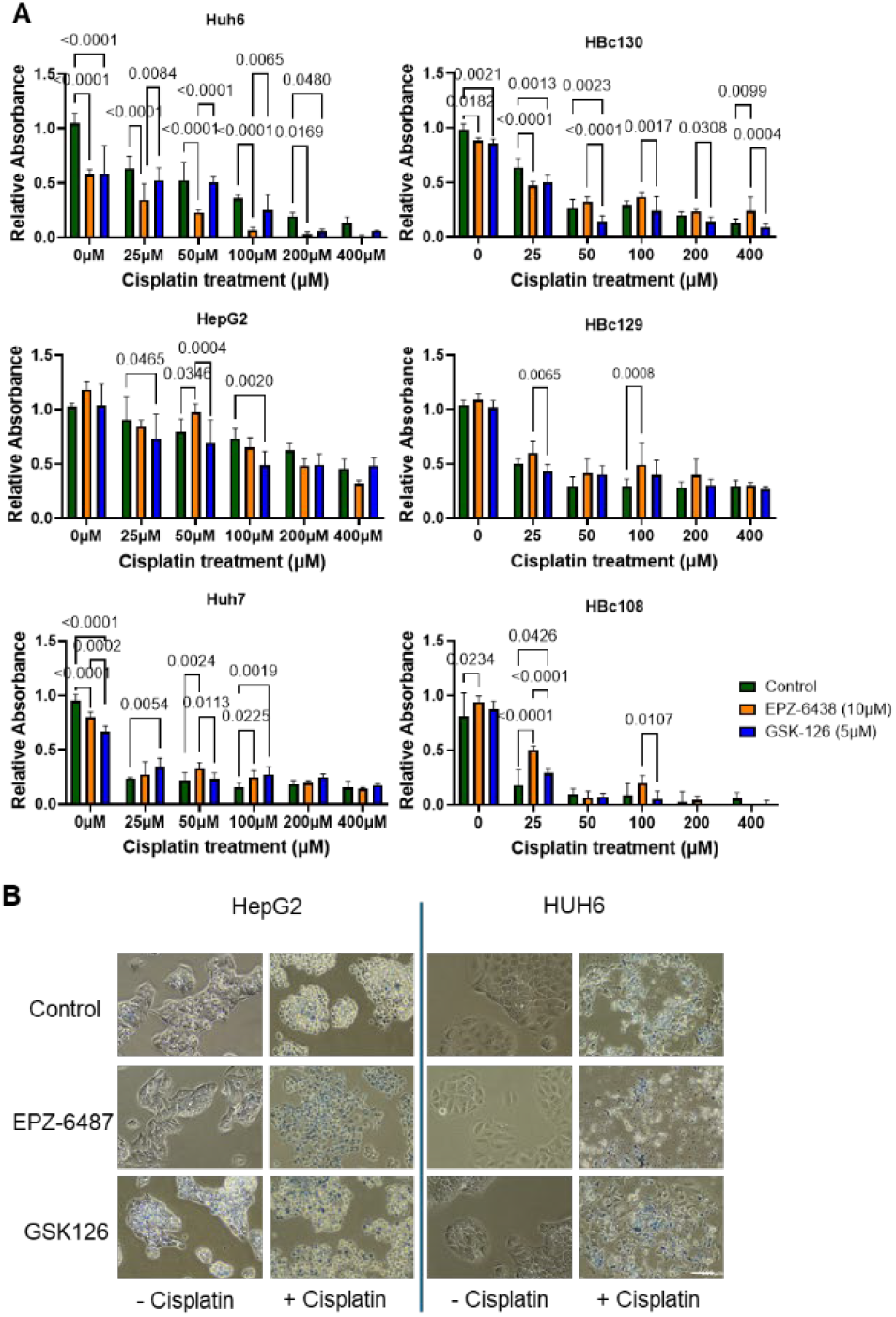
EZH2 inhibition makes HB cells more sensitive to Cisplatin treatment. A. Liver cancer cells treated with EZH2 inhibitors *in vitro* followed by cisplatin demonstrated differential response to treatment. HCN-NOS showed no response to EZH2 inhibitors alone (HBc108). HB cells, HUH6 and HBc130 showed improved response to cisplatin in combination with EZH2 inhibition, with variability between the inhibitors. HepG2 cells showed elevated cisplatin resistance but better response with EZH2 inhibition. HBc129 showed limited response to cisplatin and no added benefit of EZH2 inhibition, while HUH7, HCC cells showed better cisplatin response but no additive effect with EZH2 inhibitors. B. Cell morphology suggests improved cell death and synergy between EZH2 inhibitors and cisplatin treatment. (Images represent EZH2 inhibitors added for 24 hours, then 400μM cisplatin plus inhibitors for an additional 24 hours – 20X images with trypan blue to highlight remaining dead cells.)

Control cells exhibited normal morphology with intact cell membranes and dense cytoplasm (**Figure 5B**). EPZ-6438 and GSK-126 monotherapy induced mild morphological changes, suggesting limited cytotoxicity when used alone. Combination treatments (+cisplatin) led to morphological alterations, including cell shrinkage, detachment, and membrane blebbing—hallmarks of apoptosis.

### EZH2 inhibition reduces tumor growth and improves response to Cisplatin treatment

Treatment of one HB PDX model (HB66 model) with EPZ-6438 (150 mg/kg), cisplatin (5 mg/kg), or their combination demonstrated differential effects on tumor growth over a 28-day period (**Figure 6A, B**)(40). Monotherapy with either EPZ-6438 or cisplatin resulted in moderate tumor growth suppression compared to the untreated control group. However, the combination therapy produced a markedly enhanced reduction in tumor volume, with the most pronounced effect observed by day 21. Tumors in the combination group showed a sustained and progressive decrease in relative volume, whereas tumors in the monotherapy groups exhibited slower and less consistent responses. Notably, some animals in the no-treatment and cisplatin-only groups were sacrificed early, either due to reaching terminal tumor volume thresholds or as part of a parallel study. These findings support a synergistic effect of EZH2 inhibition and chemotherapy, highlighting the therapeutic potential of combination strategies in HB.

**Figure 6.**
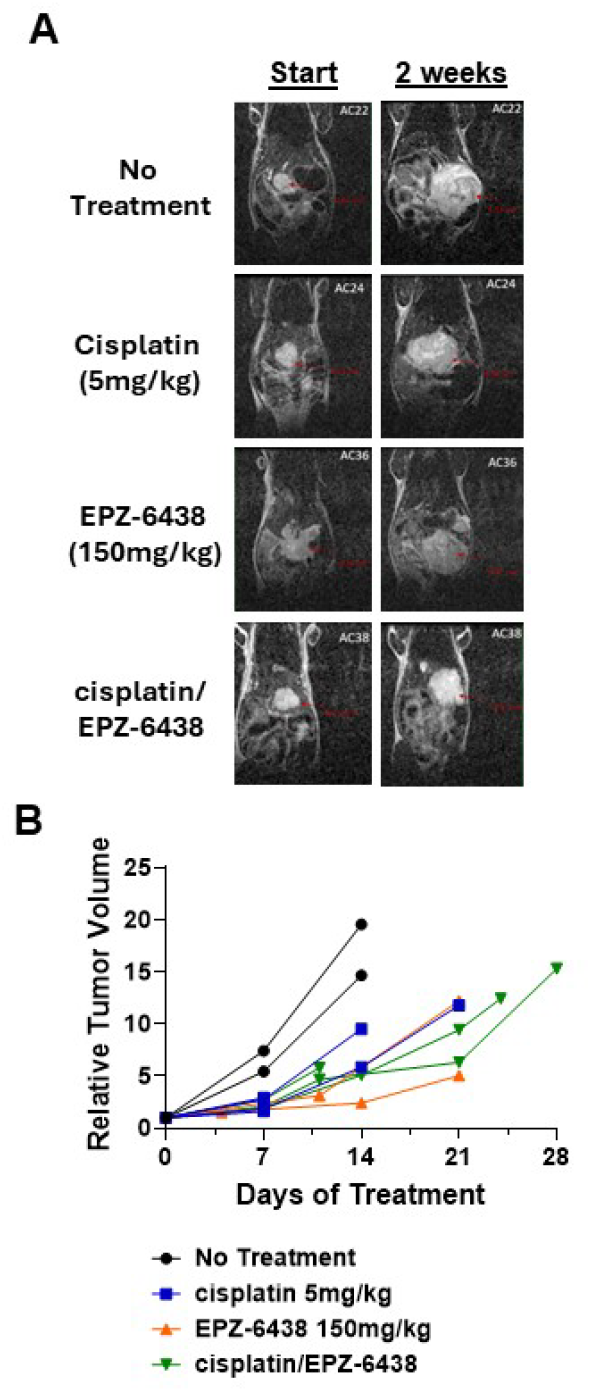
EZH2 inhibition improves *in vivo* tumor response to cisplatin in HB PDX model. HB66 PDX model was pretreated with 150mg/kg EPZ-6438 5 times per week. 5mg/kg cisplatin was then started and EPZ-6438 continued to study termination. A. MRI images show tumor volume at initiation of treatment and at 2 weeks of treatment. B. When comparing tumor volume, cisplatin alone showed modest reduction in tumor growth, while EPZ-6438 and cisplatin plus EPZ-6438 showed even greater reduction in overall tumor growth, supportive of *in vitro* studies.

### EZH2 and SUZ12 Variants in HB

In a panel of 11 HB patient tumors sequenced using the CCHMC CinCseq clinical panel, variants to both EZH2 and *SUZ12* were found in all patients (**Figure 7A**). Nine of 11 patients had variants in CTNNB1 consistent with the expected mutation range for this gene (45). All HB samples examined had c.118-4 or c.118-5 variants, with one patient also exhibiting c.553G>C and c.159G>T *EZH2* variants. *EZH2* is a key component of PRC2 which catalyzes the trimethylation of histone H3 at lysine 27 (H3K27me^3^), leading to transcriptional repression (46). All 11 samples also showed *SUZ12* c.175_177del and/or c.823+5del variants (**Figure 7B**). *SUZ12* and *EED are* other critical components of PRC2, essential for its stability and function (47). Nine of 11 patients also had EED variants. The c.118-4del variant in the *EZH2* gene is located in an intronic region, specifically four bases upstream of exon 2. This position is close to the splice acceptor site, which could potentially affect splicing. However, current evidence suggests that this deletion does not significantly impact splicing or the function of the EZH2 protein. *EZH2* and *SUZ12* variants have been reported in other cancers and can be viewed through OncoKB (48). These variants are not currently linked to clinical significance but given the overexpression of EZH2 and canonical and noncanonical functions, they may affect downstream interactions. Further work is warranted because a deletion that disrupts the splice acceptor site typically occurs within the consensus sequence at the 3’ end of an intron, just before the start of an exon. This region is crucial for proper splicing, and any alteration can lead to incorrect splicing of the mRNA (49,50).

**Figure 7.**
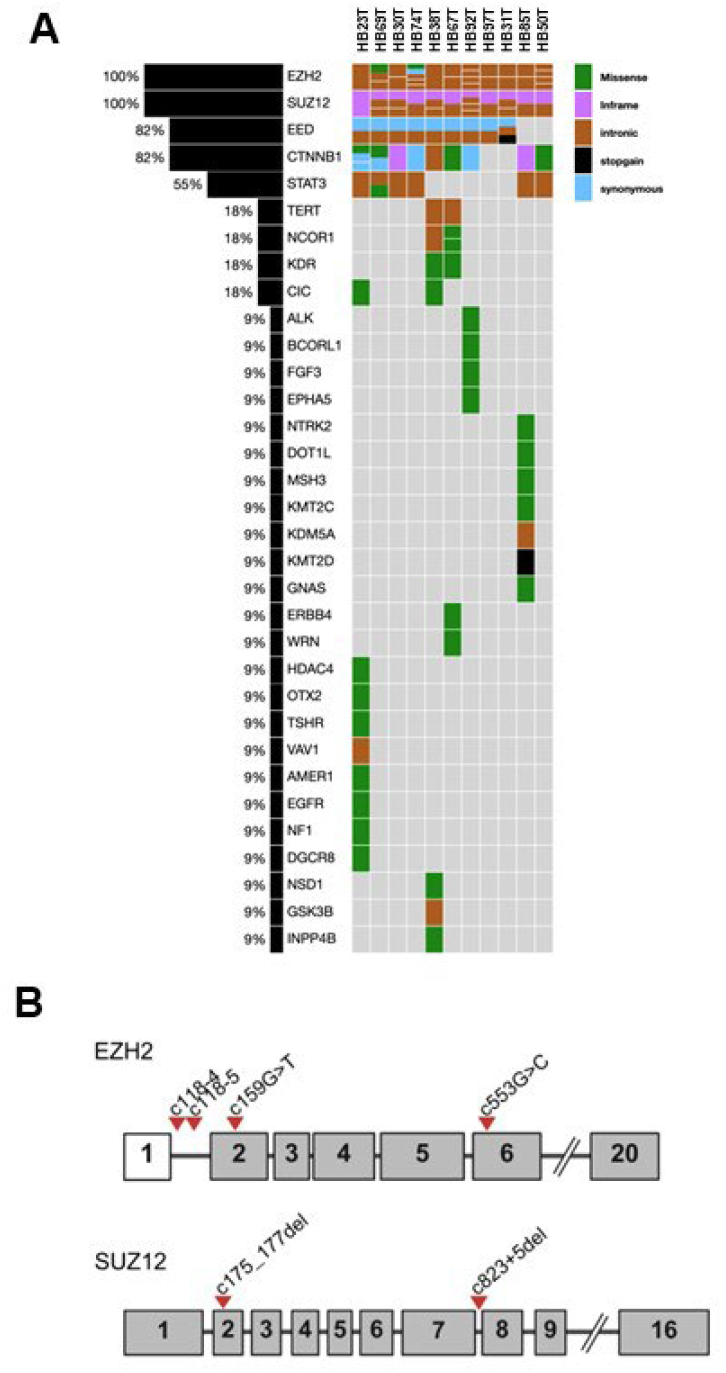
Qualitative next generation sequencing using CinCseq cancer panel to define genetic sequence variants. A. 11 patient tumors were sequenced from FFPE tissue to explore clinically relevant variants as well as VUS. EZH2 and SUZ12 have VUS in 11 out of 11 samples and EED in 9 out of 11. As expected, CTNNB1 variants are observed in 9 out of 11 patients. STAT3 variants are also observed but at a lower frequency. Other less frequently observed variants are also displayed. B. Schematic gene map drawings to represent the location of detected VUS in EZH2 and SUZ12 (all exons not shown).

## DISCUSSION

Our integrative analysis of HB using snRNA-seq revealed that EZH2 is predominantly overexpressed in cycling Hep_T_ cells, previously classified as the Tr2 subcluster. This overexpression was associated with the embryonal histologic subtype; a more aggressive clinical phenotype compared to the fetal subtype. Additionally, targeted sequencing using the CCHMC CinCseq clinical panel identified variants of uncertain significance (VUS) in both *EZH2* and *SUZ12* in a cohort of 11 HB patient tumors. Together these results highlight a complex interplay between epigenetic dysregulation, mitotic control, and histologic subtype, with EZH2 emerging as a central oncogenic driver in HB.

Unsupervised clustering and cell-type-specific marker analysis of snRNA-seq data identified distinct tumor cell subpopulations, with a cycling Hep_T_ cell population correlating with more aggressive embryonal histology. These cells exhibited disproportionate expression of PRC2 components, including EZH2, SUZ12, and EED, suggesting aberrant PRC2 signaling. Interestingly, SUZ12 was frequently localized to the cytoplasm and expressed at lower levels, indicating potential disruption of canonical PRC2 complex formation and function.

Our transcriptional profiling revealed that *EZH2* overexpression in HB is associated with upregulation of mitotic regulators such as *AURKB, Ki67*, and *CDK1*, suggesting that EZH2 promotes cell cycle progression and genomic instability. Importantly, reduced expression of CDH1, a known suppressor of WNT signaling, may further stabilize *CTNNB1*, compounding the proliferative signaling cascade in HB.

Our snRNA-seq analysis revealed transcriptional heterogeneity within HB, which aligns with previously described WNT signaling states. Consistent with the findings of Kluiver *et. al*., we observed distinct tumor subpopulations with varying levels of WNT pathway activation, which may contribute to differential drug sensitivities (43). Notably, our cycling Hep_T_ population showed strong transcriptional similarity to the embryonal population identified by Wu *et. al*., further supporting the link between proliferative capacity and WNT-driven lineage programs in HB (42). These findings underscore the relevance of WNT signaling heterogeneity in shaping both tumor biology and therapeutic response.

Functional validation using HB cell lines and PDX models demonstrated that pharmacological inhibition of EZH2 with EPZ-6438 significantly reduced proliferation and sensitized cells to cisplatin-induced cell death. This effect was most pronounced in the HUH6 cell line, where EPZ-6438 pretreatment followed by cisplatin treatment led to dose-dependent cytotoxicity within 24 hours. GSK-126 also enhanced cisplatin sensitivity, though to a lesser extent. In contrast, HepG2 and HUH7 cells, which resemble HC-NOS and HCC phenotypes, retained partial chemoresistance, suggesting tumor-type-specific mechanisms of EZH2 dependency.

Despite modest repression of canonical PRC2 targets such as CCND1 and STAT3, the discordant expression and localization of PRC2 components suggest that EZH2 may exert its oncogenic effects through noncanonical mechanisms. Of particular interest is the observed downregulation of *STAT3* at the gene level and its negative regulation in the snRNA-seq data. This could be linked to an intronic *EZH2* variant near the splice acceptor site of exon 3, potentially altering EZH2’s regulatory capacity. Previous studies have shown that EZH2 can directly methylate STAT3 and modulate its activity in a context-dependent manner. The precise role of EZH2 in STAT3 regulation in HB remains to be elucidated, however our findings suggest a novel regulatory axis that may be therapeutically exploitable.

Taken together, our data support a model in which EZH2 overexpression drives HB tumor progression through both canonical and noncanonical mechanisms, associated with dysregulated mitosis, altered PRC2 complex integrity, and potential epigenetic modulation of CDH1 signaling. The differential response to EZH2 inhibitors between HB and HCC cells further underscores the need for tumor-specific therapeutic strategies. EPZ-6438 appears to be a more effective EZH2 inhibitor than GSK-126 in HB, despite their similar mechanisms of action.

Further studies are warranted to dissect the mechanistic underpinnings of EZH2-mediated oncogenesis in HB and to evaluate the therapeutic potential of EZH2 inhibition, particularly in combination with standard chemotherapeutics. Given the association between EZH2 overexpression and embryonal histology, targeting EZH2 may offer a promising strategy to improve outcomes in this aggressive pediatric liver cancer.

## ACKNOWLEDGEMENTS

This research was supported by The Grace Foundation (AJB), Helis Foundation (SAV), Macy Easom Cancer Research Foundation (SAV), NCI R01282467 (SEW), DOD CDMRP Career Development Award W81XWH2110396 (SEW). Additionally, this project was supported by NIH P30 DK078392 Gene Analysis Core and Integrative Morphology Core of the Digestive Diseases Research Center in Cincinnati, OH and Cincinnati Children’s Hospital Research Foundation. This research was made possible, in part, using the Cincinnati Children’s Integrated Pathology Research Facility (RRID:SCR_022637), Integrated Genomics and Microbiome Sequencing Facility (RRID:SCR_022630, RRID:SCR_022653, RRID:SCR_022649), and Veterinary Services.

## Notes

The authors declare no potential conflicts of interest.

### Competing Interest Statement

The authors have declared no competing interest.

