## Supplemental Tables S1 and S2 for "Oncogenic Role of Aberrant EZH2 in Hepatoblastoma"

**Supplemental Table 1 - Patient Sample Demographics**

| Patient ID | Description | Gender | Age at sample collection (years) | Pretext stage | Histology | Predominant histologic subtype | Neoaduvant Chemotherapy prior to acquisition |
|---|---|---|---|---|---|---|---|
| 17 | HB | F | 5.21 | IV | fetal | fetal | Y |
| 18 | HB | M | 2.21 | III | mixed epithelial, fetal, embryonal, minor mesenchymal | mixed | Y |
| 21 | HB | M | 2.96 | IV | Predominant pattern pleomorphic; embryonal; HCC-like | embryonal | Y |
| 23 | HB | F | 1.51 | III | variable histology, no classic fetal/embryonal | | Y |
| 24 | HB | F | 3.81 | IV | HCC-like | | Y |
| 25 | HB | M | 2.02 | IV | cartilage | | Y |
| 27 | HB | F | 1.73 | IV | Crowded fetal, rare multinucleated and pleomorphic, blastemal, pseudoacinar | mixed | Y |
| 28 | HB | F | 1.58 | III | bone, cartilage | | Y |
| 29 | HB | M | 1.88 | IV | Metastatic HB, embryonal, undifferentiated morphology | embryonal | Y |
| 30 | HB | M | 3.40 | IV | Crowded fetal, embryonal, blastemal, rare pleomorphic | transition | Y |
| 31 | HB | M | 2.19 | II | mixed epithelial (fetal and embryonal) and mesenchymal without teratoid features | mixed | Y |
| 38 | HB | M | 2.90 | IV | Crowded fetal, blastemal, cholangioblastic | fetal | Y |
| 40 | HCC | M | 27.08 | I | Well-differentiated HCC | HCC | N |
| 41 | HB | M | 2.06 | IV | Crowded fetal, embryonal, blastemal | mixed | Y |
| 42 | HB | M | 3.09 | IV | HCC-like (pleomorphic, steatosis) | | Y |
| 43 | HB | M | 1.51 | I | epithelial type, fetal, blastemal | fetal | N |
| 45 | HB | M | 7.98 | III | Epithelial type, macrotrabecular, embryonal, fetal | mixed | Y |
| 46 | HB | F | 2.64 | II | mixed fetal and embryonal, mesenchymal type without teratoid features | mixed | Y |
| 47 | HB | M | 3.59 | III | epithelial type, fetal (crowded), embryonal, blastemal, pleomorphic | mixed | Y |
| 48 | HCN-NOS | M | 11.90 | IV | fetal-like, macro, mild pleomorphism | HCN-NOS | Y |
| 49 | HB | M | 0.38 | III | epithelial type, fetal, embryonal (minor amount), blastemal, pleomorphic | fetal | Y |
| 50 | HB | F | 0.64 | II | Crowded fetal, blastemal, osteoid | fetal | Y |
| 52 | HCN-NOS | M | 12.15 | IV | HCN-NOS | HCN-NOS | Y |
| 53 | HB | F | 2.20 | IV | Crowded fetal, embryonal, blastemal | embryonal | Y |
| 60 | HB | M | 3.57 | II | Epithelial type, predominantly embryonal with fetal and blastemal components | embryonal | Y |
| 62 | HB | | | | Metastatic HBL, mesenchymal, osteoid | mesenchymal | Y |
| 64 | HB | M | 1.63 | IV | Metastatic HBL, mesenchymal, osteoid | mesenchymal | Y |
| 66 | HB | F | 3.94 | III | epithelial type, mixed fetal and embryonal with focal macrotrabecular and minor blastemal | mixed | Y |
| 67 | HB | F | 5.04 | II | epithelial type, mixed embryonal and fetal blastemal, pleomorphic | mixed | Y |
| 69 | HB | F | 0.84 | III | Crowded fetal | fetal | Y |
| 70 | HB | M | 4.00 | II | Metastatic HB, epithelial type, predominantly embryonal with crowded-fetal and minimal blastemal component | embryonal | Y |
| 71 | HCN-NOS | M | 7.56 | IV | HCN-NOS | HCN-NOS | Y |
| 73 | HB | M | 1.19 | IV | epithelial, predominantly fetal, only small embryonal | fetal | Y |

| | | | | | | | |
|---|---|---|---|---|---|---|---|
| 74 | HB | F | 3.35 | IV | Crowded fetal, embryonal, blastemal, mild pleomorphic | embryonal | Y |
| 75 | HCN-NOS | M | 16.00 | III | HCN-NOS | HCN-NOS | Y |
| 76 | HB | F | 1.08 | III | fetal | fetal | Y |
| 77 | HCN-NOS | M | 13.64 | IV | HCN-NOS | HCN-NOS | Y |
| 79 | HCC | F | 16.44 | IV | HCC | HCC | Y |
| 80 | HB | F | 0.63 | II | Epithelial, fetal | fetal | Y |
| 81 | HB | M | 4.65 | II | Crowded fetal, embryonal | embryonal | Y |
| 82 | HB | F | 3.70 | IV | Pleomorphic, HCC-like areas | | Y |
| 83 | HB | F | 1.88 | III | epithelial type, embryonal, pleomorphic (poorly differentiated), SCUD, mesenchymal type without teratoid features | mixed | Y |
| 84 | HCC | M | 13.20 | I | HCC, moderately differentiated | HCC | N |
| 85 | HB | M | 6.59 | I | epithelial type, embryonal, fetal (crowded), cholangioblastic and blastemal, focal mesenchymal | mixed | Y |
| 86 | HB | M | 3.29 | | Metastatic HB; blastemal with focal mesenchymal present | | Y |
| 87 | HB | F | 1.31 | III | mixed epithelial and mesenchymal, fetal pattern (mitotically inactive), mesenchymal without teratoid | fetal | Y |
| 91 | HB | F | 3.02 | II | mixed epithelial and mesenchymal, fetal (mitotically active), embryonal, pleomorphic (poorly differentiated) | mixed | Y |
| 92 | HB | F | 0.82 | II | epithelial type, fetal (mitotically active), blastemal, with cholangioblastic differentiation | | Y |
| 94 | HB | F | 0.99 | IV | epithelial type, fetal pattern, mesenchymal | fetal | Y |
| 96 | UES | F | 6.64 | II | undifferentiated embryonal sarcoma | UES | Y |
| 97 | HB | F | 2.83 | IV | fetal | Fetal | Y |
| 99 | UES | F | 6.91 | II | undifferentiated embryonal sarcoma | UES | Y |
| 105 | HCC | M | 10.96 | | moderately differentiated | HCC | N |
| 108 | HCN-NOS | M | 11.50 | III | Pleomorphic fetal of HB overlap with HCN-NOS | fetal | Y |
| 129 | HB | M | 4.43 | II | mixed fetal, crowded fetal and blastemal with focal pleomorphic and cholangioblastic differentiation | mixed | Y |
| 130 | HB | M | 5.20 | I | crowded fetal, embryonal, mixested with blastemal | mixed | Y |

### Supplemental Table 2- Antibodies and real time PCR primers

| Antibodies | Dilution | Company | Product Number | Publication |
|---|---|---|---|---|
| *IHC/IF* | | | | |
| EZH2 (IHC, IP-Western) | 1:200 | Cell Signaling | 3147 | PMID: 35301492 |
| SUZ12 | 1:1,000 | Cell Signaling | 3737 | PMID: 36612203 |
| EED | 1:200 | Cell Signaling | 85322 | PMID: 36428492 |
| Alexa Fluor 488 goat anti-mouse IgG | 1:2,000 | Invitrogen | A11001 | |
| Alexa Fluor 555 goat anti-rabbit IgG | 1:2,000 | Invitrogen | A21428 | |

| Gene Primers | Company | Product Number |
|---|---|---|
| EZH2 | Qiagen | PPH02880A |
| SUZ12 | Qiagen | PPH17208A |
| EED | Qiagen | PPH23422 |
| CTNNB1 | Qiagen | PPH00643F |
| Ki67 | Qiagen | PPH01024E |
| GAPDH | Qiagen | PPH00150F |
| AURKB | Qiagen | PPH21059F |
| GPC3 | Qiagen | PPH11457B |
| STAT3 | Qiagen | PPH00708F |
| CDH1 | Qiagen | PPH00135F |
| TGFβ | Qiagen | PPH00508A |
| MYC | Qiagen | PPH00100B |
